# Mosquitoes reared in distinct insectaries within an institution in close spatial proximity possess significantly divergent microbiomes

**DOI:** 10.1101/2024.08.28.610121

**Authors:** Laura E. Brettell, Ananya F. Hoque, Tara S. Joseph, Vishaal Dhokiya, Emily A. Hornett, Grant L. Hughes, Eva Heinz

**Affiliations:** Department of Vector biology, Liverpool School of Tropical Medicine, Liverpool, L3 5QA, UK; School of Science, Engineering and Environment, University of Salford, Manchester, M5 4WT, UK; The Roslin Institute, Royal (Dick) School of Veterinary Studies, The University of Edinburgh, Midlothian, EH25 9RG, UK; Department of Evolution, Ecology and Behaviour, University of Liverpool, Liverpool, L69 7ZB, UK; Department of Tropical Disease Biology, Liverpool School of Tropical Medicine, Liverpool, L3 5QA, UK; Department of Clinical Sciences, Liverpool School of Tropical Medicine, Liverpool, L3 5QA, UK; Strathclyde Institute of Pharmacy and Biomedical Sciences, University of Strathclyde, G4 0RE, Glasgow, UK

## Abstract

The microbiome affects important aspects of mosquito biology and differences in microbial composition can affect the outcomes of laboratory studies. To determine how the biotic and abiotic conditions in an insectary affect the composition of the bacterial microbiome of mosquitoes we reared mosquitoes from a single cohort of eggs from one genetically homogeneous inbred *Aedes aegypti* colony, which were split into three batches, and transferred to each of three different insectaries located within the Liverpool School of Tropical Medicine. Using three replicate trays per insectary, we assessed and compared the bacterial microbiome composition as mosquitoes developed from these eggs. We also characterised the microbiome of the mosquitoes’ food sources, measured environmental conditions over time in each climate-controlled insectary, and recorded development and survival of mosquitoes. While mosquito development was overall similar between all three insectaries, we saw differences in microbiome composition between mosquitoes from each insectary. Furthermore, bacterial input via food sources, potentially followed by selective pressure of temperature stability and range, did affect the microbiome composition. At both adult and larval stages, specific members of the mosquito microbiome were associated with particular insectaries; and the insectary with less stable and cooler conditions resulted in slower pupation rate and higher diversity of the larval microbiome. Tray and cage effects were also seen in all insectaries, with different bacterial taxa implicated between insectaries. These results highlight the necessity of considering the variability and effects of different microbiome composition even in experiments carried out in a laboratory environment starting with eggs from one batch; and highlights the impact of even minor inconsistencies in rearing conditions due to variation of temperature and humidity.

## Introduction

The microbiome profoundly affects diverse aspects of mosquito biology. It is critical for larval development and influences survival, reproduction and immunity (Cansado-Utrilla et al., 2021; Martinson & Strand, 2021; Salgado et al., 2024). The microbiome can furthermore impact the transmission of pathogens by mosquitoes; either indirectly by impacting mosqutio life span or reproduction, or directly by interfering with or facilitating pathogen establishment in the host (Cansado-Utrilla et al., 2021; Hughes et al., 2014). Indeed, microbial-based control strategies are proving to be successful avenues for vector control (Ross et al., 2022). However, our understanding of both how the microbiome affects the mosquito host, and how its assembly as a complex community takes place, is far from complete.

The composition of the mosquito microbiome can vary substantially depending on a range of biotic and abiotic factors. The microbiomes of field-caught mosquitoes are affected by host species, geography and local climate (Bascuñán et al., 2018; Hegde et al., 2018; Jeffries et al., 2024; Medeiros et al., 2021). Laboratory-reared mosquitoes commonly used for experimental studies, on the other hand, harbour a simpler microbiome, and mosquitoes respond differently to these microbiomes of differing complexities (Hegde, Brettell, et al., 2024; Santos et al., 2023). It has become apparent that despite the relative stability of the insectary environment, microbiome differences can be seen between both species, and between genetically homogenous and inbred mosquito lines (i.e., the same species derived from different field-collected individuals) under the same rearing conditions (Coon et al., 2014; Kozlova et al., 2021; Saab et al., 2020).

Laboratory studies using *Aedes aegypti*, the major vector of arboviruses including dengue, Zika and yellow fever viruses have shown variations in the microbiome between generations, and when transferred to new institutions (Accoti et al., 2023; Saab et al., 2020). Conversely, another study found mosquitoes from diverse geographic origins reared in a common insectary environment harboured remarkably similar microbiomes (Dickson et al., 2018). Taken together, these results strongly suggest the local insectary environment or rearing conditions affect microbiome composition. This perhaps is unsurprising, since bacteria are readily taken up by mosquitoes through feeding as larvae and adults (Coon et al., 2022; Kulkarni et al., 2021; MacLeod et al., 2021). However, other studies have reported different *Ae. aegypti* lines, reared in the same insectary environment, show differences in their microbiome composition demonstrating the role of the host in microbiome selection (Kozlova et al., 2021; Short et al., 2017).

Given the complex reciprocal interactions, it can be challenging to disentangle the role of the host, the environment (e.g. larval water) and abiotic conditions (e.g. temperature) on host-associated microbiome composition. In human disease research, a ‘reproducibility crisis’ has implicated the gut microbiome as a critical determinant of the reproducibility and translatability of research performed using animal models (Dirnagl et al., 2022). In particular, work with laboratory-reared mice with the same genetic background has found strong facility effects on the microbiome (Parker et al., 2018). This has resulted in researchers recommending the reporting or consideration of microbiome composition in studies using laboratory mice (Ericsson & Franklin, 2021). Similarly, elucidating these interactions in mosquitoes has implications for interpreting results of laboratory-based studies, in particular considering the impact the microbiome can have on pathogen transmission, which have notoriously been variable (Bennett et al., 2002; Gubler & Rosen, 1976; Kilpatrick et al., 2010; Roundy et al., 2017; Tesh et al., 1976) .

To understand the influence of the insectary environment on the mosquito microbiome without the confounding effects of host genetics and potential vertically transmitted microbiome components, we reared mosquitoes from a single cohort of *Ae. aegypti* eggs in three different insectaries and characterised their bacterial microbiome composition at both the larval and adult life stages. Complementary to this we assessed the microbiome composition of input food sources used for rearing, recorded environmental conditions within the insectaries, and noted host development times and survival rates. Our work furthers the understanding of the relative influence that host and environment exert on the microbiome composition in mosquitoes. We conclude that it is important to understand and characterise the mosquito microbiome for the accurate evaluation of laboratory studies using mosquitoes.

## Methods

### Experimental Setup

The study took place across three different insectaries (here called A, B and C), within 200m of each other at the Liverpool School of Tropical Medicine (LSTM) (Figure 1a). All insectaries are within 200 m of each other. All insectaries are regularly used by multiple research groups to maintain long term mosquito lines and to carry out mosquito experiments. During the experiment, insectary A also housed colonies of *Anopheles gambiae*, *Anopheles stephensi*, *Aedes albopictus* and additional *Ae. aegypti* lines. Insectary B housed a colony of *Culex pipiens* and there were no other mosquitoes in insectary C. The insectaries resource fish food from the same provider. The three insectaries’ conditions were set according to standard user protocols of 27 °C / 75% relative humidity (RH) (insectary A), 25 °C / 60% RH (insectary B) and 26 °C / 75% RH (insectary C) (Supplementary Table 1). The three insectaries were set at different set conditions were to allow for a favourable environment for the specific mosquito species housed there, with insectary B being commonly used to rear temperate mosquito species and insectaries A and C being used for tropical/subtropical species. To monitor temperature (°C) and relative humidity (%), a Tinytag Ultra 2 data logger (Gemini data loggers, UK) was placed within each insectary, next to larval trays, recording every 15 minutes for the duration of the experiment. Whilst the insectaries are within the same institution, they are in buildings differing in age. Insectary A - 2007, B - 1903/1904 (refurbished 2010/2012) and C - 2017.

**Figure 1:**
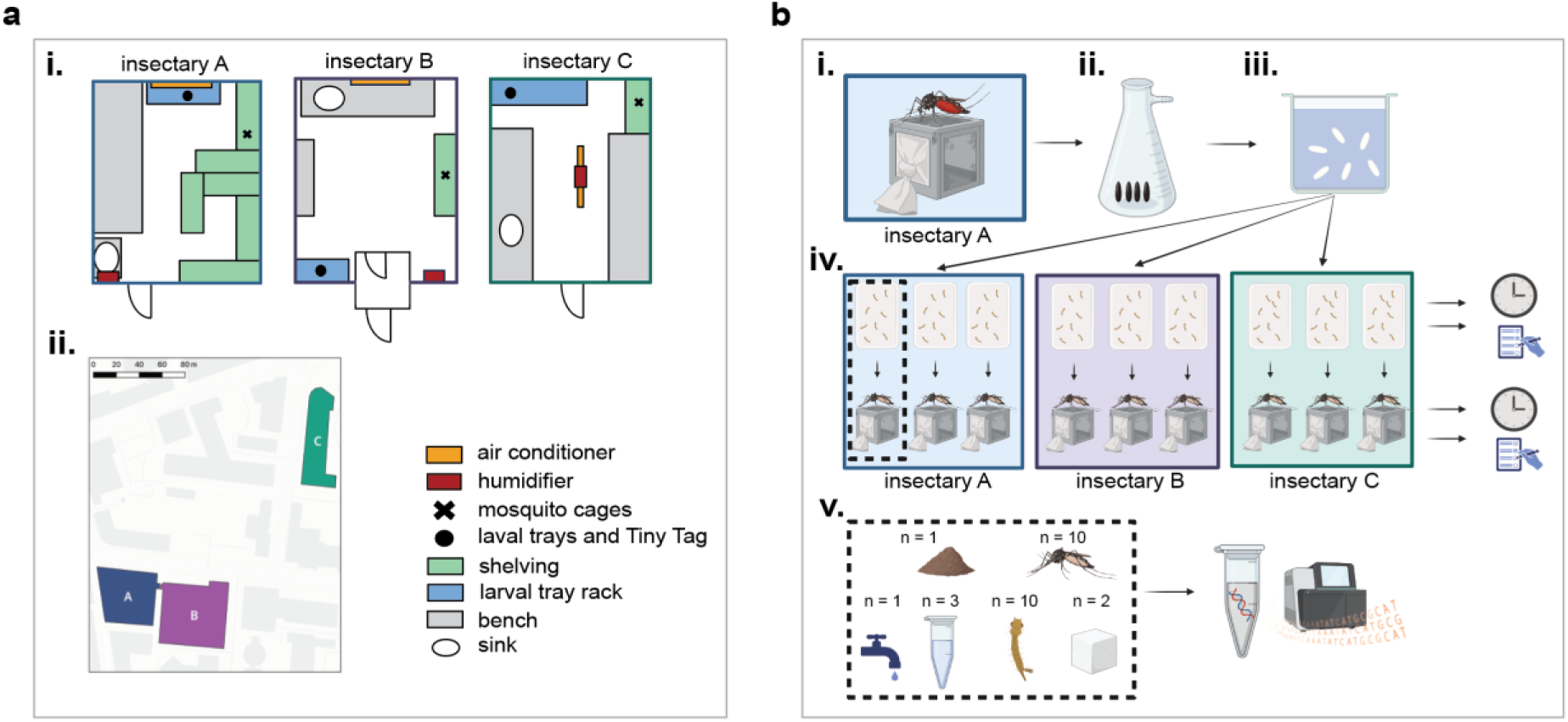
Layout of the insectaries used in this experiment and experimental setup. a: Schematic showing the layouts of each individual insectary used in this experiment, with **i.**) placement locations of mosquito trays and cages and **ii.**) map showing locations of the three buildings where insectaries are located. **b:** Experimental setup. **i.)** Conventionally reared *Ae. aegytpi* (Liverpool line) that had been continually reared in ‘insectary A’ at the Liverpool School of Tropical Medicine (LSTM) were allowed to lay eggs under standard conditions. **ii.)** One cohort of eggs were vacuum hatched in the laboratory. **Iii.)** The resulting L1 larvae were divided into nine trays of 150 larvae. **iv.)** Three replicate trays were transferred into each of three insectaries at LSTM: the original insectary ‘insectary A’, and two further insectaries ‘insectary B’ and ‘insectary C’. Here, the cohorts were reared to adulthood according to standard conditions, recording the number of individuals that successfully developed to pupal and adult life stages. Recordings were always made between 09:00 and 12:00. TinyTag data loggers were used to measure the temperature and humidity throughout the experiment. **v.)** For each of the three replicates in each of the three insectaries (shown in dashed line box), the following samples were collected: one fish food sample, one tap water sample, three larval water samples and ten L3/L4 larvae samples collected at the same time, two sugar solution samples and ten adult females. One additional tap water sample was also collected from each insectary. Samples were then stored at -80 °C, before **vi.)** DNA extraction along with an additional extraction blank per batch and 16S rRNA sequencing. Panel **a ii**. was created with QGIS version: version 3.28, https://www.gqis.org/ Basemap: Positron, Map tiles by CartoDB, under CC BY 3.0. Data by OpenStreetMap, under ODbL. Panel **b** was created with Biorender.com.

A cohort of eggs was derived from a single colony of *Ae. aegypti* reared in insectary A (Figure 1b). The mosquitoes belonged to the ‘Liverpool line’ that are descendants of an original west African colony brought in to the laboratory in 1936 and which are continually maintained at LSTM (Ramachandran et al., 1960). The colony used to generate eggs for this study included 300-400 adult females which were provided with fresh human blood from the National Health Service before being provided with moist filter paper to lay eggs. The resulting egg paper was dried before splitting into small segments which were randomly assigned to three equal batches. These segments were vacuum hatched for 45 minutes in tap water sourced from each respective insectary. The hatched larvae were then transferred to the three insectaries, fed with one spoon (approx. 0.3 g) of TetraMin fish food (Tetra), and placed in a larval tray with 1 L tap water overnight to develop. The tap water and fish food were obtained from each insectary’s own taps/stocks, with the fish food from insectaries A and B originating from one batch and the fish food from insectary C from another. Trays were cleaned between uses with hot soapy water and were kept in each insectary, with insectaries A and B routinely sharing trays. Four replicate samples of tap water (2 ml per sample) and three of fish food (0.3 g per sample) were collected per insectary for microbiome analysis and stored at -80 °C. The following day, larvae in each insectary were further split into three new replicate trays per insectary with 150 larvae per tray. Each tray was fed with 0.3 g of fish food every two days and monitored daily for survival. Pupation began on day 7, at which point pupae from each tray were picked and transferred to a small container of fresh tap water within a corresponding cage. Pupae were picked for 3 days in total between 09:00-12:00, after which the number of larvae which had failed to develop were recorded. Each cage of adults was provided with sugar solution (10% sucrose) throughout the experiment. Sugar solution is routinely prepared by combining table sugar with distilled water in a glass bottle that has been cleaned with hot soapy water. Distilled water was obtained from the nearest available source, which is the same for insectaries A and B and different for insectary C. Stocks of sugar solution are stored on a benchtop in each insectary and replenished once empty. Ten individual larvae were collected from each tray when they reached L3/L4 stage, along with three replicate samples of larval water per tray (2 ml per tray). Numbers of hatched adults were counted on day 14. Ten adult females were collected from each cage at 3-5 days post-emergence (days 12-14) and two replicate sugar water samples (2 ml per sample) were collected per cage. Larvae and adult mosquitos were surface sterilised in 70% ethanol, then washed and stored in sterile 1X PBS. All samples were frozen at -80 °C until processed.

### DNA extraction and library preparation

Genomic DNA from all samples was extracted using Qiagen DNA Blood and Tissue kit with modified protocols. For insect tissue (whole adults and larvae), samples were homogenized in sterile 1X phosphate-buffered saline (PBS) and incubated with 80 µl proteinase K and 180 µl ATL lysis buffer for 3 hours at 56 °C. The remaining extraction steps were performed following the manufacturer’s supplementary protocol for DNA extraction from insect cells. Water (both tap water and larval water) and sugar samples (10% sucrose) were first centrifuged at 8000 rpm for 10 minutes. Then, the supernatant was removed and pellets were resuspended in 180 µl enzymatic lysis buffer (containing 20mM Tris-Cl (pH 8.0), 2mM sodium EDTA, 1.2% Triton X-100 and 20 mg/ml lysozyme) and incubated for 30 minutes at 37°C. Samples were then incubated with 25 µl proteinase K and 200 µl buffer AL at 56°C for 30 minutes, before continuing the subsequent steps from the manufacturer’s instructions. For fish food samples, 2 ml sterile 1X PBS was added to each 0.3 g sample and vortexed to obtain a homogenous mixture. Samples were then centrifuged at 8000 rpm for 10 minutes and the pellet was subjected to DNA extraction following the above protocols. A blank extraction control (extraction process used for water and sugar samples, but with sterile water as input) was included with each batch of DNA extractions (n = 7) to account for extraction or kit contaminants.

DNA was quantified using fluorometry (Qubit) and shipped on dry ice to Novogene, Cambridge, UK, for library preparation using primers targeting the hypervariable V4 region of the 16S ribosomal RNA gene (515F and 806R (Caporaso et al., 2011)) and sequencing on the Novaseq 6000 to generate 250bp paired end reads.

### Data analysis

Raw sequence reads (fastq format) were denoised using DADA2 (Callahan et al., 2016) and taxonomy was assigned to amplicon sequence variants (ASVs) by applying the classify-sklearn algorithm in QIIME 2 (v2022.2) using a Naïve Bayes classifier pre-trained on the SILVA 138.1 database (Quast et al., 2012). The phylogenetic relationships between ASVs were determined in QIIME 2 through a multiple sequence alignment using MAFFT (Katoh & Standley, 2013) and phylogenetic reconstruction using fasttree (Price et al., 2009). QIIME data artifact (qza) files were then imported into Rstudio ((R Core Team, 2023); v4.3.2) for subsequent analyses. These data were then converted to a *Phyloseq* object (McMurdie & Holmes, 2013) and the *Decontam* package (Davis et al., 2018) was then used to identify and remove contaminant ASVs using the ‘prevalence’ method and following recommendations from (Díaz et al., 2021) to identify contaminants as all sequences more prevalent in controls than true samples. The dataset was then filtered further to remove mitochondria and chloroplast sequences and retain only bacterial ASVs using the subset_taxa command in the *Phyloseq* package. Rarefaction curves were generated for all samples, with the exclusion of the negative controls, remaining after quality control and filtering using the ‘ggrare’ function in the *Ranacapa* package (Kandlikar et al., 2018), followed by rarefaction at the smallest library size (post filtering). The resulting rarefied counts table was then used for all subsequent analyses.

Alpha (Shannon’s index) diversity was calculated using the *MicrobiotaProcess* package (Xu et al., 2023) and plotted using *ggplot2* (Wickham, 2011). Statistical significance in between groups were calculated using Kruskal Wallace Rank Sum tests using the ‘kruskal.test’ function in the *stats* package v4.3.2 (R Core Team, 2023) with *post hoc* pairwise testing using Dunn’s tests with Bonferroni adjustment for pairwise testing (Dinno, 2017). Differences were considered statistically significant if *p* ≤ alpha/2. Beta diversity metrics (Bray-Curtis and unweighted Unifrac) were calculated using the *Phyloseq* package with the ‘distance’ function, followed by ordination using the ‘ordinate’ function and plotting using ‘plot_ordination’. Ellipses were added to the plots using ‘stat_ellipse’ using the default 95% confidence levels assuming multivariate t-distribution. Overall differences in beta diversity between sample types were calculated using permutational multivariate analysis of variance (PERMANOVA) with the ‘adonis2’ function in the *vegan* package (Oksanen J, 2022), with subsequent pairwise comparisons calculated using the ‘pairwise.adonis2’ function in the *pairwiseAdonis* package (Arbizu, 2017). Differences between groups were considered statistically significant if *p* ≤ 0.05. To identify whether there were statistically significant differences between samples from the different insectaries, data were subset by sample type and distance metrics recalculated. For each sample type, ‘adonis2’ and ‘pairwise.adonis’ tests were again used to determine whether samples from the three insectaries were statistically significant. For the larvae, larval water and adult female samples, adonis2 was also used to determine whether there were cage/tray effects by assessing the nested interaction of try/cage within insectary. Relative abundance plots were created from the *Phloseq* object, with *ggplot2.* Determination of differentially abundant bacteria between the three insectaries was carried out with the ‘ancombc2’ function in the *ANCOM* package (Lin & Peddada, 2020, 2024). Multiple pairwise comparisons between each insectary was carried out using a fixed formula of insectary + sample type and controlling the overall mdFDR at 0.05 using the Holm-Bonferroni method. Heatmaps showing relative abundance of ASVs were generated using the ‘plot_heatmap’ function in the *Phyloseq* package.

Numbers of individuals successfully developing to pupal and adult stages in each replicate tray/cage were recorded at days two and nine respectively and visualised using *ggplot2* with differences between insectaries calculated using Kruskal-Wallis tests using the kruskal.test function in Rstudio (v4.3.2). Time to pupation was also recorded for each replicate tray and plotted. At the completion of the experiment, insectary condition measurements (temperature and relative humidity) were downloaded from the TinyTag data loggers in csv format. Minimum, maximum and mean temperatures were calculated for each insectary and plotted in Rstudio (v4.3.2) using *ggplot*. Brown-Forsythe tests were then used to test for differences in spread of the data between the three insectaries using the ‘bf.test’ function in the *onewaytests* package (Dag et al., 2018).

Scripts for all analyses and figure generation are available at https://github.com/laura-brettell/insectary_comparison.

## Results

### Abiotic environmental factors and mosquito development show differences between insectaries

Temperature and humidity differed between the three insectaries across the experiment. While slight differences were to be expected due to different research groups’ protocols requiring slightly different set values (Supplementary Table 1), we also observed marked differences in their deviations from set values (Figure 2 a, b, Supplementary table 2). Fluctuations within each insectary correlated between temperature and relative humidity. Insectary A experienced the most variable temperature (av. = 27.81 °C, std dev = 1.78), with some days on average 4.49 °C higher than others. Insectary B, on the other hand, experienced the most variable humidity (av = 51.8 %, std dev = 3.72). Insectary C was notably more consistent than the other insectaries, with minimal variations to temperature (av = 26.31 °C, std dev = 0.12) and humidity (av = 80.00 %, std dev = 0.60). Insectary A, the most highly used of the three, showed notable differences over the course of the experiment and insectary B showed most variable conditions each day and a decrease in fluctuations in the last four days of the experiment. We noted no major change in frequency or mode of use in any of the insectaries over the duration of our experiment with the exception of reduced activity during weekends.

**Figure 2:**
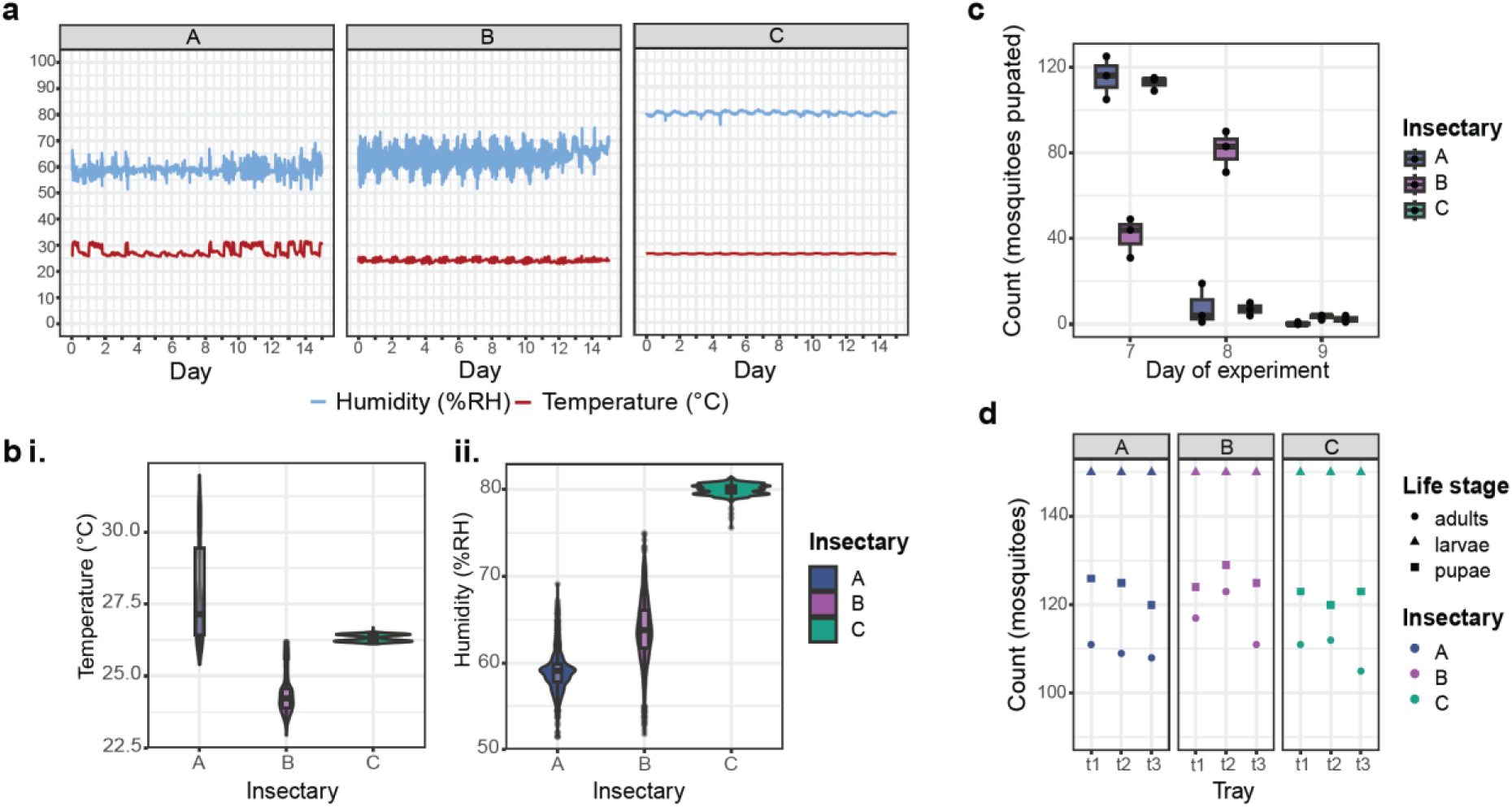
Environmental conditions and mosquito development in each insectary over the course of the experiment. a: Temperature (°C) and humidity (%RH) were recorded every 15 minutes using TinyTag data loggers in insectaries A, B and C. Weekends were days five/six and 12/13 and there were no public holidays during this time. **b**: Average and spread of recorded temperature (**i.**) and humidity (**iii.**) in each insectary. **c**: Time taken for individuals to develop to the pupal stage in each insectary. **d**: Mosquito development in each replicate tray, faceted by insectary, showing numbers of individuals successfully developed to the pupal and adult stages from an initial 150 larvae/tray.

Mosquito development was monitored in the three insectaries over 14 days and showed no statistically significant difference in the numbers of mosquitoes that successfully developed to pupal and adulthood life stages in each insectary (Figure 2d, Supplementary Table 3). We note more variation between trays and less uniform and longer development times in insectary B (Figure 2c) which is also the insectary with the lowest temperature, and strongest daily fluctuations in temperature and humidity (Figure 2a, b).

### Microbiome complexity varies in mosquitoes reared in different insectaries and in their food sources

Altogether, 16S rRNA amplicon sequencing was carried out for 253 samples comprising 90 adult females, 90 L3 larvae, 27 larval water samples, 18 sugar solution samples, 12 tap water samples, nine fish food samples and seven extraction blanks (Figure 1a). After quality control and filtering, 244 samples remained, comprising 89 adult females, 89 L3 larvae, 27 larval water samples, 18 sugar solution samples, 12 tap water samples and nine fish food samples. These generated an average of 43,907 reads per sample (ranging from 3,974 to 74,250) (Supplementary Table 4). Samples were then rarefied to the lowest sampling of 3,974 reads/sample, at which point the majority of rarefaction curves had plateaued (Supplementary Figure 1).

Overall, alpha diversity (Shannon’s Index) was significantly different between sample types (Kruskal-Wallis, ꭕ^2^ = 65.93, *p* = <0.001). To account for these distinct profiles per sample type, pairwise differences in alpha diversity between insectaries were compared for each sample type separately. Both larvae and larval water samples showed statistically significant pairwise differences between those from insectary B and those from both insectaries A (larvae: Dunn’s test, *z* = -6.56, *p* = <0.001 and larval water: *z* = -4.72, *p* = <0.001) and C (larvae: *z* = 4.32, *p* = <0.001 and larval water: *z* = 2.41, *p* = 0.024), with samples from insectary B showing the highest alpha diversity (Figure 3a, Supplementary Table 5). Conversely, adult mosquitoes showed no statistically significant differences in alpha diversity between insectaries. While the sugar solution samples were significantly different in alpha diversity between insectaries B and C (*z* = 3.30, *p* = 0.002), with insectary B exhibiting a lower diversity. There were no differences in alpha diversity of the tap water or fish food samples between any insectaries. However, the fish food samples from insectaries A and B, which originated from the same batch, were observably more diverse than the fish food from insectary C which originated from a different batch. These samples, comprising amongst other ingredients fish and crustacean derivatives, yeasts and algae, appeared highly variable both within and between insectaries. We do acknowledge, that this dried material might also contain a substantial amount of DNA remnants from bacteria that were present in fish and other components.

**Figure 3:**
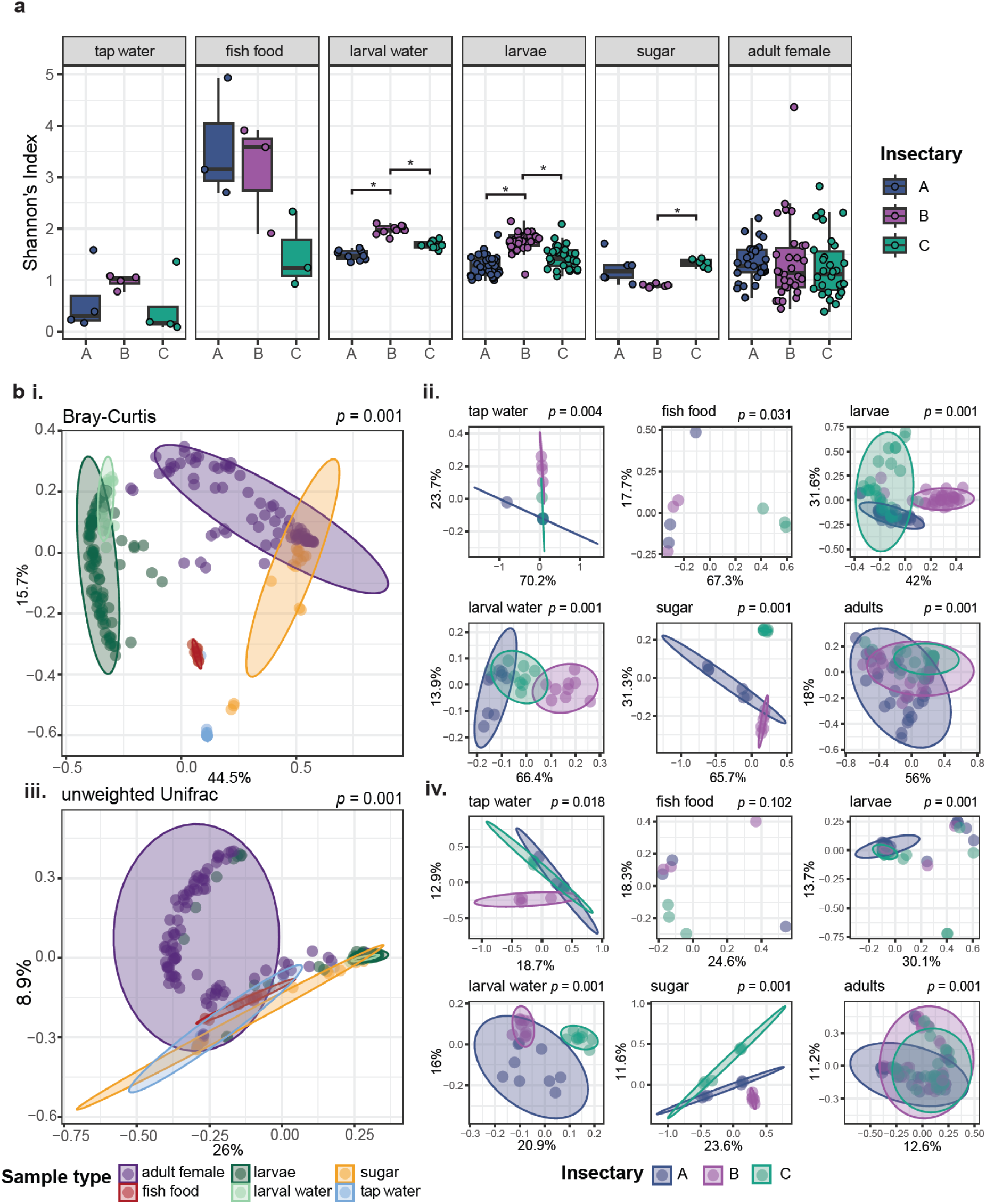
**Microbial diversity amongst sample types from different insectaries**. **a**) Alpha diversity calculated as Shannon’s index for each sample type, grouped by insectary (A, B, C). Statistically significant pairwise differences between samples from the three different insectaries, within sample types, are denoted by asterisks and are calculated using Kruskal Wallace tests with post-hoc pairwise Dunn tests (*p* value ≤ alpha/2). **b**) PCoA plots showing beta diversity calculated as (**i, ii**) Bray-Curtis and (**iii, iv**) unweighted Unifrac dissimilarity metrics. Diversity was calculated using all samples passing quality thresholds, and coloured according to sample type (**i, iii**). Diversity metrics were then recalculated on the data subset by sample type and coloured to visualise distribution of samples originating from each of the three insectaries (**ii, iv**). *p* values show results of PERMANOVA analyses to determine differences between sample types (**i, iii**) insectary within each sample type (**ii, iv**).

**Figure 4:**
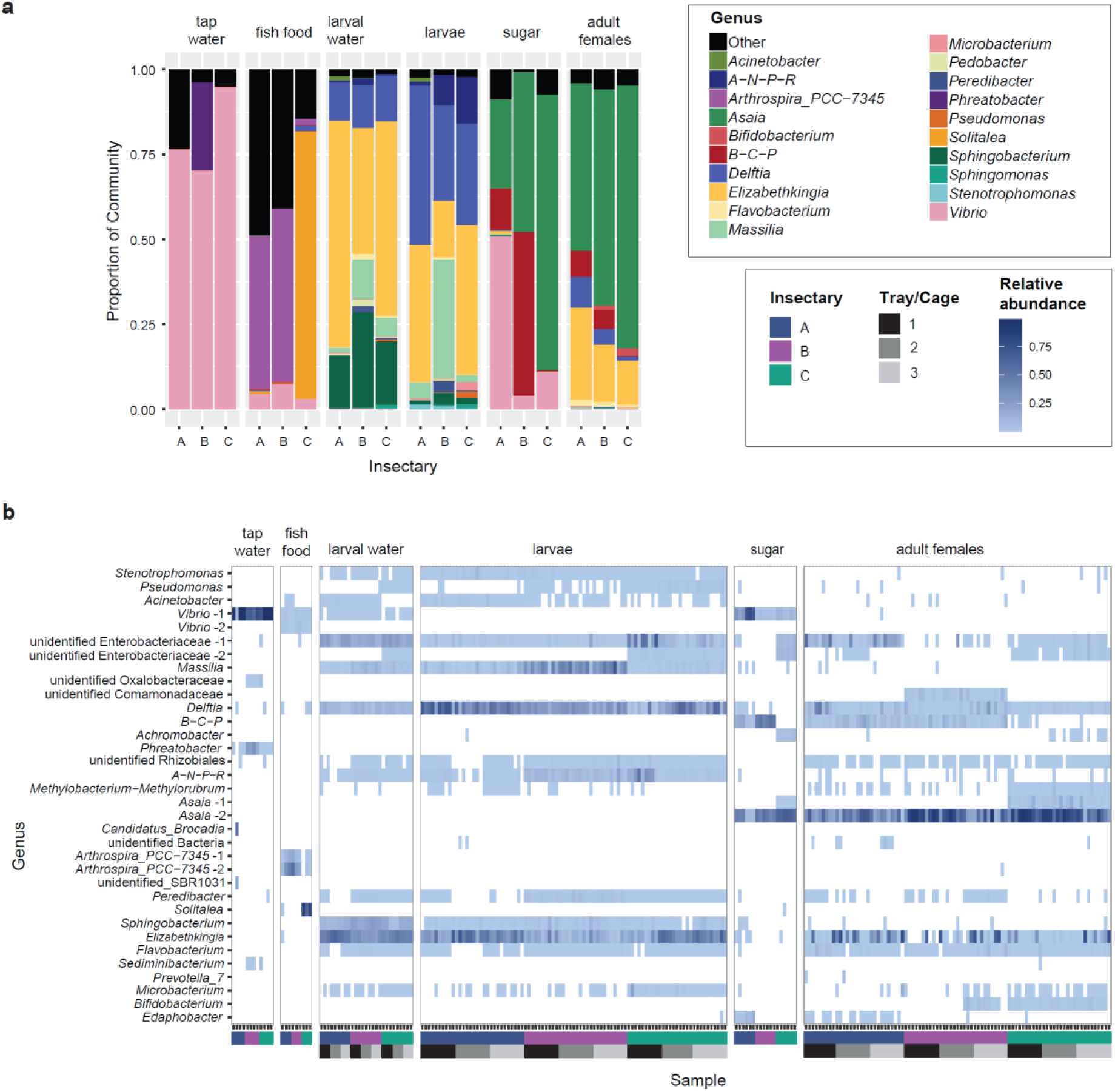
**Taxonomic composition of the microbiome across ample types and insectaries**. **a**) Relative abundance of the top 20 most abundant genera in the data set averaged according to whether they were from insectary A, B or C, for each sample type (tap water, fish food, larval water, larvae, sugar and adult females). All other genera were grouped together as ‘Other’. Detailed per-sample composition is shown in Figure S2. **b)**: Heat map showing the relative abundance of ASVs in each sample, including all ASVs present at ≥ 5% relative abundance in at least one sample. Each row corresponds to a single ASV and is labelled on the y axis according to genus if known or, if unknown, the lowest taxonomic ranking known. Where there are taxonomic groups containing more than one ASV present at ≥ 5% relative abundance in at least one sample, the labels are suffixed with a number (eg ‘*Asaia* - 1’). Each column corresponds to a single sample, faceted by sample type. Upper colour blocks on the x axis denote insectary of origin. Lower colour blocks denote tray/cage number within each insectary for larval water, larvae and adult female samples. Tap water, fish food and sugar samples were collected before being provided to trays/cages. Relative abundance is indicated by the blue gradient, with more highly abundant ASVs in darker shade. Zero values are indicated in white.

Beta diversity analysis showed statistically significant differences between each sample type using both Bray-Curtis and unweighted Unifrac distance metrics (adonis *p* ≤ 0.005, Figure 3b i, ii, Supplementary Table 6). For each sample type, there were also significant differences between insectaries using both metrics, with the exception of the fish food samples using unweighted Unifrac dissimilarity (Figure 3b ii, iii, Supplementary Table 7). Furthermore, larvae, larval water and adult female samples all showed statistically significant cage/tray effects, using both metrics (Supplementary Table 7).

### Compositional microbiome differences in food and at larval stages converge during mosquito development

Given differences in diversity, we next assessed the taxonomic composition of the dataset for differences between different sample types and insectaries. As expected, following our observations on similarities in beta diversity, there were clear similarities in identified taxa between samples of the same sample types (Figure 3a, b, Supplementary Figure 2). Considering the composition of different sample types averaged within an insectary, adult female mosquitoes were dominated by *Asaia* and *Elizabethkingia*. Larvae and larval water samples were similar in composition and dominated by *Delftia* and *Elizabethkingia*, with *Delftia* also detected in adult mosquitoes from all insectaries, and the larval water also contained a high proportion of *Sphingobacterium* ASVs. Tap water samples were dominated by *Vibrio* and these were also present in the sugar and fish food samples albeit at lower abundances, but not present in larval or adult mosquito samples (relative abundance < 0.00). Sugar samples from all insectaries also contained a high proportion of *Asaia* sequences. The fish food samples for all three insectaries contained dominant genera not seen in other sample types, and that varied between insectaries. The fish food from insectary C was dominated by *Solitalea* (78.6%), which used a different fish food stock to insectaries A and B, which were dominated by Arthrospira_PCC-7345.

While the sample types contained a similar composition of main taxa in the three insectaries, the relative abundances of these genera varied by insectary (Figure 3a), and across individual samples (Supplementary figure 2). Across the data averaged by sample type, in the larval samples there were strong differences between *Massilia* (4.7, 35.1 and 19.5%, in insectaries A, B, C, respectively) and *Elizabethkingia* (40.5, 16.6 and 44.1%); and *Asaia* varied in sugar samples between 26.3, 47.2 and 81.2%. Adult mosquitoes showed differences mainly in the ratio of *Asaia* (49.2, 63.7 and 77.4%) and *Elizabethkingia* (27.2, 16.9 and 12.9%), and a smaller but varying distribution of *Delftia* (9.0, 4.6 and 1.3%) and *Burkholderia-Cabelleronia- Paraburkholderia* (7.8, 5.5 and 0.1%). Within sample types, we observed individual variation, which appeared to be greatest in the adult females (Supplementary figure 2). Despite the clear differences between sample types, there were bacteria that showed statistically significant differences between insectaries across the dataset as a whole (Supplementary Figure 3). Most notably *Burkholderia-Cabelleronia-Paraburkholderia* was more abundant in insectaries A and B than insectary C (Ancom-bc, log fold changes of 3.82 and 4.02 respectively, Supplementary Figure 3).

Following the detection of cage/tray effects in beta diversity, we used Ancom-bc to assess whether particular taxa were differently abundant in samples from different trays (larvae and larval water samples) and cages (adult females) in the different insectaries. Tap water, sugar and fish food were not assessed as these were collected prior to providing to a tray/cage. Differentially abundant taxa were seen between trays and cages in all insectaries, however the majority of differentially abundant taxa were specific to one insectary and either trays or cages (Supplementary figure 4). Only *Delftia* was identified as differentially abundant in all three insectaries (between cages in insectary A and trays in insectaries B and C). The differentially abundant bacteria comprised both dominant bacteria in the relevant sample types, including *Massilia* which was differentially abundant between trays in insectary C, and bacteria which were present at much lower abundances including *Stenotrophomonas* which was differentially abundant between cages in insectary B.

To assess differential composition at higher resolution, we assessed whether different ASVs from the same genus, which can indicate different species or lineages, were present associated with insectaries and potentially restricted to specific trays/cages. Whilst the majority of dominant genera were only represented by one ASV, some of the dominant genera, including *Asaia* and *Vibrio,* comprised multiple ASVs, which may represent different species/lineages with different biological functions (Figure 3b). Further indicating insectary- specific microbiomes, specific ASVs were present in different sample types from the same insectary not apparent at the genus level. Most notably, one *Asaia* ASV (“Asaia 1”) was present in adult female and sugar samples from insectary C, but this was not present in samples from either of the other insectaries. Further, one ASV within the Enterobacteriaceae was common in samples from insectary C (“unclassified Enterobacteriaceae 2”), and present in the majority of mosquito samples across all life stages. However, this ASV was far less common in insectary A, only detected in 10/29 adult females, and absent from insectary B samples.

## Discussion

To understand how the insectary environment can affect microbiome composition whilst controlling for host background, we used a single cohort of *Ae. aegypti* eggs, split into three batches, and reared these in three different insectaries in parallel. Microbiomes can be affected by a range of external and host factors, so we measured key environmental parameters as well as assessed microbial diversity of potential input sources (tap water, fish food, larval water, sugar solution). We then recorded mosquito development and monitored the establishment of the microbiome in larvae and adult female mosquitoes.

The microbial diversity between the different insectaries was comparatively similar when considering the main taxa per sample type, with the exception of fish food. Mosquito microbiomes were dominated by bacterial genera commonly seen in mosquito studies, including *Asaia*, *Elizabethkingia* and *Delftia* (Foo et al., 2023; Lin et al., 2021; Scolari et al., 2019). Differences in bacterial input via food sources affected microbiome composition in the different insectaries. This was particularly apparent in the adult stage, where *Asaia* was a dominant genus in both the mosquito microbiomes and the sugar water on which they fed. One *Asaia* ASV was present in samples from all insectaries, whereas a second was present only in the sugar and adult mosquitoes from one insectary, supporting environmental acquisition of *Asaia* from the sugar feed. This highlights how different bacterial input may be available in different insectaries and when provided with the required conditions, in this case *Asaia* being provided with sugar solution, it may become a dominant member of the mosquito microbiome. Given that members of *Asaia* have been found to exert complex interactions with *Wolbachia* and pathogens (Hughes et al., 2014; Ilbeigi Khamseh Nejad et al., 2024; Osuna et al., 2023), this illustrates the relevance to consider potential microbial variation when conducting laboratory experiments.

Taxa observed in the input samples (tap water, fish food, larval water, sugar solution) were however only selectively present in larval and adult samples, with several dominant taxa not becoming established in the mosquito microbiomes despite representing a large proportion of the input samples. Whilst the microbiome composition in fish food was different between insectaries, neither *Solitalea* nor *Arthrospira*, the two dominant taxa, were detected in the larvae or adult mosquito samples. Furthermore, the tap water, sugar and fish food samples all contained *Vibrio*, which however was absent from mosquito samples suggesting it is common in the insectary but is unable to successfully colonize the larval or persist in the adult stages, at least not to a detectable abundance, potentially due to exclusionary competition via other members of the microbiome (Hegde et al., 2018). Furthermore, the physical conditions of the mosquito provide different selection pressures that favour different bacteria to those most successful in external environments, and different species and lines of mosquitoes can vary in how they control and interact with their microbiomes (Accoti et al., 2023; Muturi et al., 2016).

At the larval stage, mosquitoes varied in their microbiome diversity between the three insectaries, with individuals from insectary B being more diverse than those from insectaries A and C. This pattern was mirrored in the larval water, with which the mosquitoes regularly exchange microbes as they develop. This is of interest given the high variance of the conditions (temperature, humidity) in insectary B, which might further drive a less stable microbiome. While we saw no statistically significant differences between the alpha diversity of adult mosquito microbiomes in the different insectaries, we did see specific ASV signatures associated with particular insectaries. One *Asaia* ASV was found in all adults reared in insectary C, but in none of those reared in insectaries A or B. Whilst we discovered *Delftia* and *Asaia* co-occurring in the same individual adult females, previous studies indicated a potential co-exclusion of *Delftia* and *Asaia* (da Silva et al., 2022). However, especially given our ASV analysis demonstrated different *Asaia* ASVs in different insectaries, it remains to be determined whether this putative negative correlation is species- or strain-specific and might thus differ between studies if only observed at 16S rRNA level. As 16S rRNA analysis cannot give insights into genetic determinants, it might be specific genome elements not present in all members of these genera that underpin the mechanisms responsible for causing co-exclusion.

Additionally, one Enterobacteriaceae ASV was present in the majority of adults, larvae and larval water from insectary C, in approximately one third of adults from insectary A, but not larvae or larval water, and was absent from samples reared in insectary B. Members of the Enterobacteriaceae can have various impacts on mosquitoes, including phenotypic effects (Dickson et al., 2017), interaction with arboviruses (Apte-Deshpande et al., 2014; Wu et al., 2019) and other bacteria in the microbiome (Kozlova et al., 2021). Furthermore, Enterobacteriaceae exposure as larvae has been shown to influence adult phenotypes (Dickson et al., 2017). Thus, different Enterobacteriaceae might have profound impacts on subsequent experiments and our data highlights the variability even in this controlled experiment with minimal influences besides the standard rearing protocol.

The biotic and abiotic conditions also differed between the three insectaries with food sources (fish food and sugar solution) differing in microbiome composition, and environmental conditions (temperature and humidity) varying in their means and variability over time. Temperature affects diverse mosquito traits such as development, fecundity and vector competence and can affect the composition of the microbiome, including across the temperature ranges seen across our study (Mordecai et al., 2019; Onyango et al., 2020; Villena et al., 2022). The effects of humidity are less well studied, in part due to the covariance with temperature and rainfall in the field, however it is also known to affect facets of mosquito biology such as egg production and desiccation tolerance (Brown et al., 2023). Instability in temperature and humidity, including diurnal shifts can also affect mosquitoes, including factors related to vector competence (Carrington, Armijos, Lambrechts, & Scott, 2013; Lambrechts et al., 2011; Pathak et al., 2024). Insectary C was remarkably stable in temperature and humidity compared to the other two insectaries, whilst the others showed a more varied pattern between and within days and larger deviations from the mean. In contrast to a previous study, we saw slower pupation times in an insectary with higher temperature fluctuations (insectary B) (Carrington, Armijos, Lambrechts, Barker, et al., 2013). Although as insectary B was also the coolest insectary, this highlights the complexity of disentangling interacting effects of means and variation in temperature, and indeed biotic factors as the larvae in insectary B harboured the most diverse microbiomes. In addition, we observed significant differences between trays and cages in all insectaries, highlighting that ideally results should try to combine mosquitoes from multiple trays to account for this, which might be driven by position in the room (especially in relation to airflow), adjacency to other species being reared, or stochastic variation of microbes associated with individual eggs which then would get transferred into the larval water.

We appreciate not all factors could be controlled here, and might have additional impact on our results. That includes potential differences in the air flow in different insectaries, and the placement of the trays and cages in relation to that which is driven practically by the spatial layout of the room. There could further be differences between cleaning regimes and disinfection methods, which we could not fully control as these are shared insectaries between multiple research groups with different experiments; a very common situation when working in research insectaries, which might impact the microbiomes. We were also not aware how the presence of other mosquito lines could impact the rearing of mosquitoes, development times or microbes present in the insectary that might get circulated in the airflow. In addition, we acknowledge the limitation of relying on 16S rRNA sequence data, which can also be derived from remnants of dead bacteria, and of only considering bacteria in the microbiome, where fungi, single-cell eukaryotes and viruses might have further impacts (Hegde, Khanipov, et al., 2024) .

## Conclusions

Laboratory experiments are commonly performed to assess diverse facets of mosquito biology under standard conditions. Whilst factors including mosquito species and line are commonly accounted for, the microbiome can also affect experimental results and is itself influenced by diverse factors. By rearing batches of *Ae. aegypti* from a single egg cohort in three insectaries at one institution, we found insectary-specific differences in microbiome diversity in mosquito larvae and adult females and specific ASVs associated with different insectaries and cages/trays. Our results highlight that rearing protocols, in particular bacterial input from food sources combined with differences in the abiotic environment likely lead to compositional changes to the mosquito microbiome.

## Data availability

All sequence reads are publicly available at Sequence Read Archive (SRA) under project code PRNJ1115112 and detailed accession numbers per sample are given in Table S4. All code used for analysis and to generate figures is available at https://github.com/laura-brettell/insectary_comparison.

## Acknowledgements

We thank staff and students from the Vector Biology Department at the Liverpool School of Tropical Medicine for generously providing insectary space for this study. This work was supported by the Biotechnology and Biological Sciences Research Council (BBSRC; BB/V011278/1, to EH and GLH) and the National Institutes of Health (NIH; R21AI138074 to GLH). GLH was further supported by BBSRC (BB/T001240/1, BB/X018024/1, and BB/W018446/1), the UK Research and Innovation (UKRI; 20197 and 85336), the Engineering and Physical Sciences Research Council (EPSRC; V043811/1), a Royal Society Wolfson Fellowship (RSWF\R1\180013), the National Institute for Health and Care Research (NIHR2000907), and the Bill and Melinda Gates Foundation (INV-048598). EH was further supported by the Wellcome Trust (217303/Z/19/Z). LEB was supported by the Liverpool School of Tropical Medicine Director’s Catalyst Fund. VD was supported by the UKRI Medical Research Council (MRC; MR/N013514/1). *The funders had no role in study design, data collection and analysis, decision to publish, or preparation of the manuscript*.

## Author contributions following CRediT taxonomy

Conceptualization – LEB, GLH, EH

Data Curation – LEB, TSJ, AFH, VD

Formal Analysis – LEB, EAH, VD,

EH Funding Acquisition – GLH, EH

Investigation – TSJ, AFH, VD

Methodology – LEB, TSJ, AFH, VD, EAH, GLH, EH

Project Administration – LEB, TSJ, AFH, VD, GLH,

EH Resources – GLH, EH

Software – LEB, TSJ, VD, EAH

Supervision – LEB, GLH, EH

Validation – LEB, TSJ, AFH, VD, EAH, GLH, EH

Visualization – LEB, TJ, VD, EH

Writing – Original Draft Preparation – LEB, AFH, TSJ

Writing – Review & Editing - LEB, TSJ, AFH, VD, EAH, GLH, EH

All authors read and approved the final manuscript version.

## Supplementary Information

**Supplementary Figure 1:**
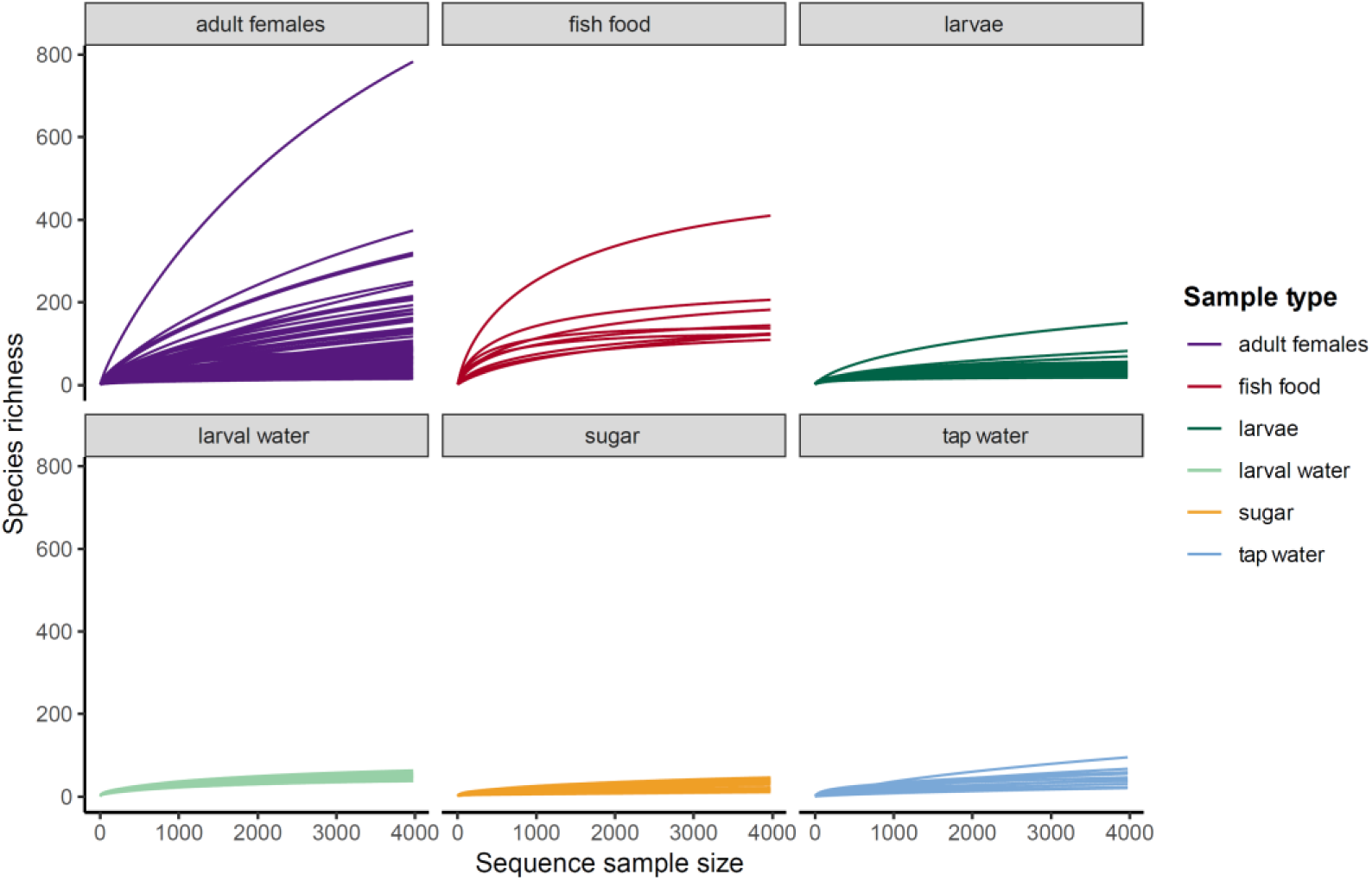
Rarefaction curves showing a plateauing for each sample type at the rarefaction depth of 3974 reads.

**Supplementary Figure 2:**
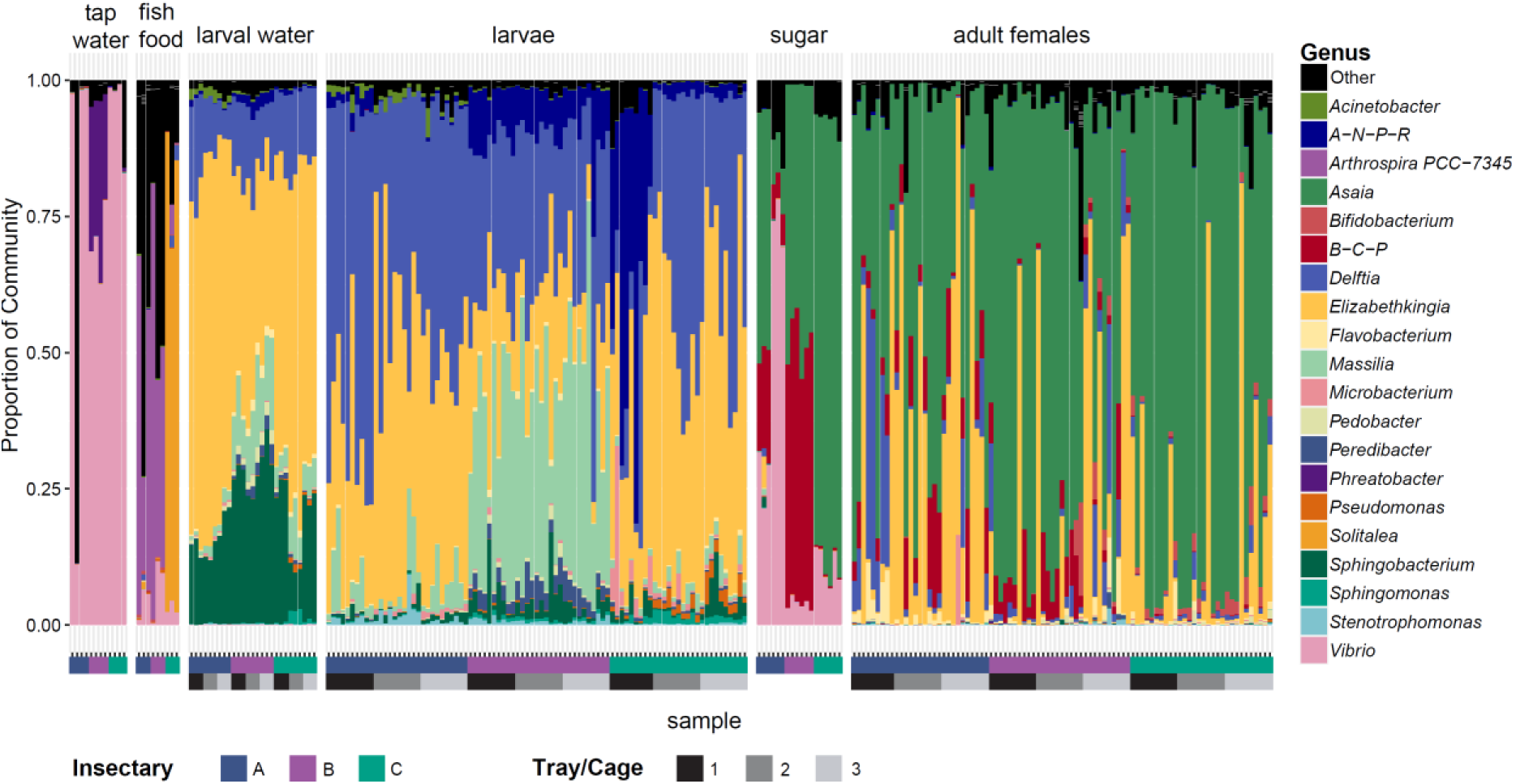
Relative abundance of the top 20 most abundant genera in the data set shown for individual samples, faceted by sample type. *Allorhizobium-Neorhizobium-Pararhizobium-Rhizobium* is abbreviated to *A-N-P-R* and *Burkholderia-Caballeronia-Paraburkholderia* to *B-C-P*. All other genera were grouped together as ‘Other’. Upper colour blocks on the x axis denote insectary of origin. Lower colour blocks denote tray/cage number within each insectary for larval water, larvae and adult female samples. Tap water, fish food and sugar samples were collected before being provided to trays/cages.

**Supplementary Figure 3:**
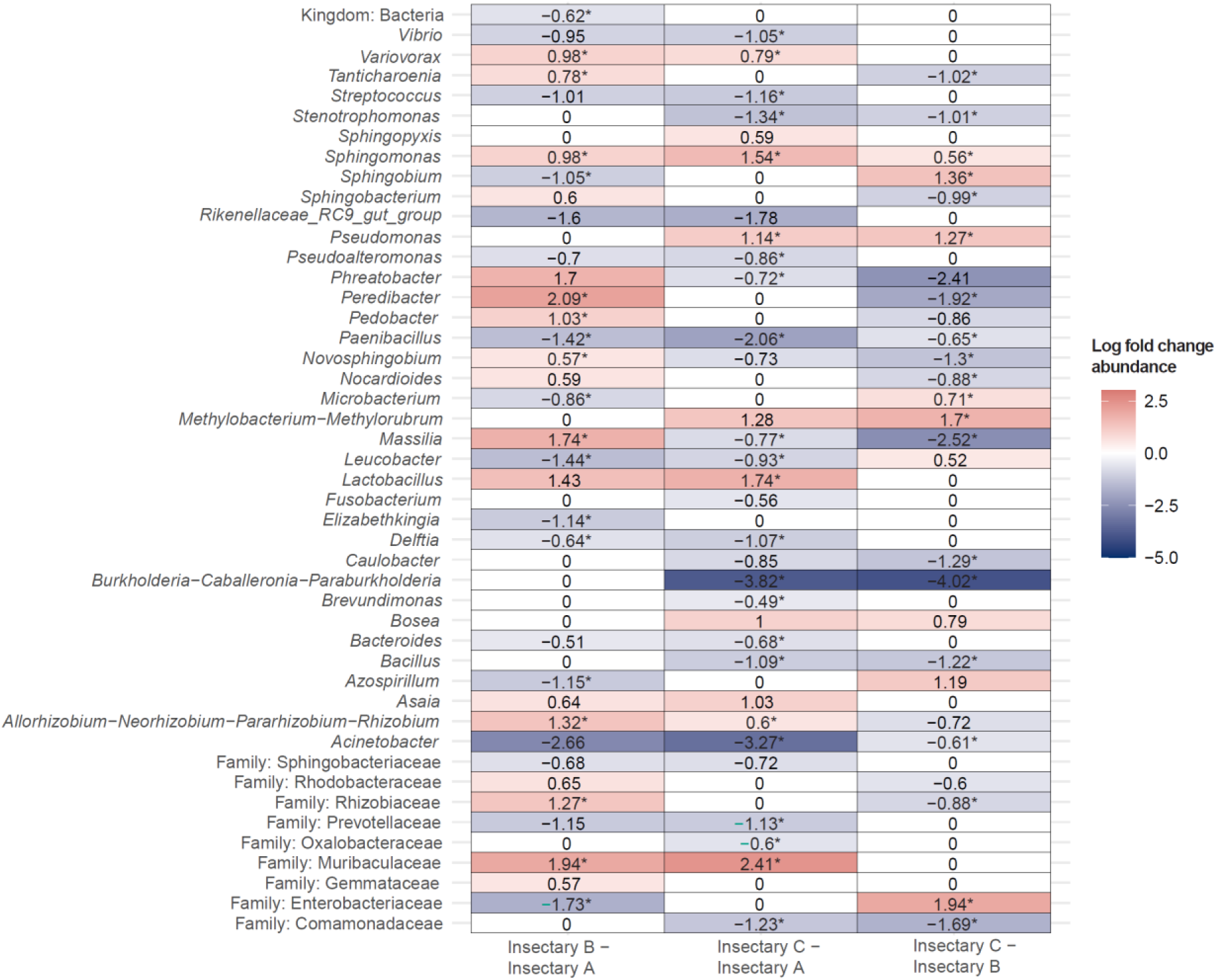
**Heatmap showing differentially abundant bacteria between each of the three insectaries in pairwise analyses**. Log fold changes are shown for each bacterial taxa, giving the highest taxonomic rank identified, which were identified as differentially abundant between insectaries (*y* axis) using ANCOM-BC2. Columns denote pairwise comparisons (i.e., column one shows log fold change in insectary B compared to insectary A) and cell colour denotes log fold change in abundance with red representing an increase in abundance and blue a decrease. Numbers represent significant changes (adjusted *p* value ≤ 0.05) and those with asterisks are significant following a further threshold of application of a sensitivity analysis for pseudo-count addition (ss filter).

**Supplementary Figure 4:**
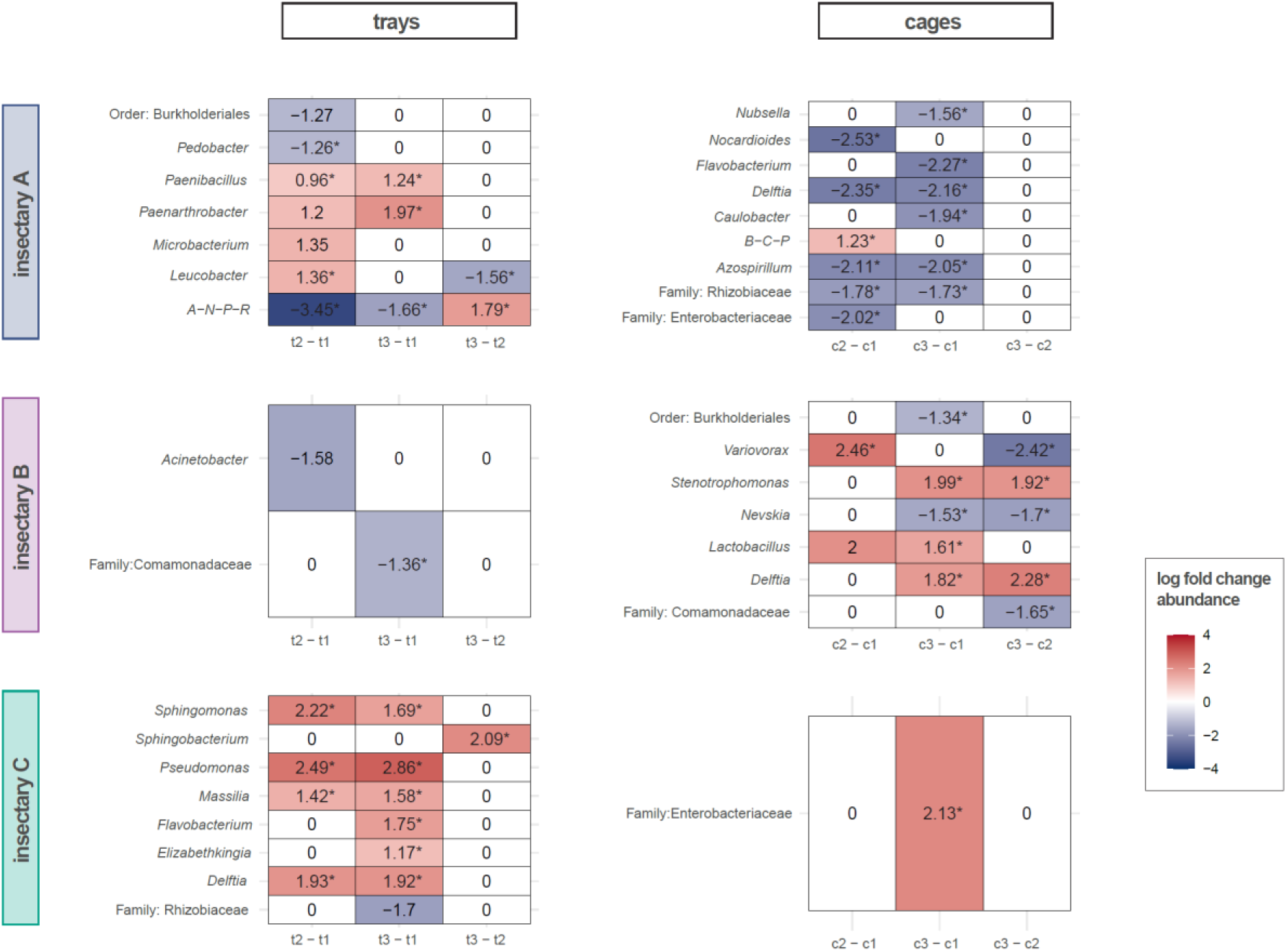
**Heatmap showing differentially abundant bacteria between trays and cages in the three insectaries, in pairwise analyses**. Log fold changes are shown for each bacterial taxa, giving the highest taxonomic rank identified, identified as differentially abundant between trays (left hand side and cages (right hand side) from insectaries A, B and C (top to bottom) using ANCOM-BC2. Rows show bacterial taxa and columns denote pairwise comparisons between trays (t1, t2, t3) or cages (c1, c2, c3) and cell colour denotes log fold change in abundance with red representing an increase in abundance and blue a decrease. Numbers represent significant changes (adjusted *p* value ≤ 0.05) and those with asterisks are significant following a further threshold of application of a sensitivity analysis for pseudo-count addition (ss filter).

**Supplementary Table 1**: Temperature, humidity and light cycle settings for the three test insectaries and average daily recorded temperature and relative humidity data using the Tinytag data logger.

**Supplementary Table 2**: Raw temperature and humidity data obtained from TinyTag data loggers.

**Supplementary Table 3**: Development data for each insectary showing numbers of mosquitoes developing to pupal and adult stages and duration to pupation.

**Supplementary Table 4**: Sample metadata for all samples passing quality control and filtering, including the sample type, building (one experimental insectary per building was used), cage/tray as applicable, the number of reads after removal of contaminant ASVs and the accession number where the raw reads can be found on Sequence Read Archive.

**Supplementary Table 5:** Results of statistical analyses of alpha diversity data.

**Supplementary Table 6**: Results of statistical analyses of beta diversity data

