## Supplementary figures and images for "Mosquitoes reared in distinct insectaries within an institution in close spatial proximity possess significantly divergent microbiomes"

### Supp_fig_1.pdf

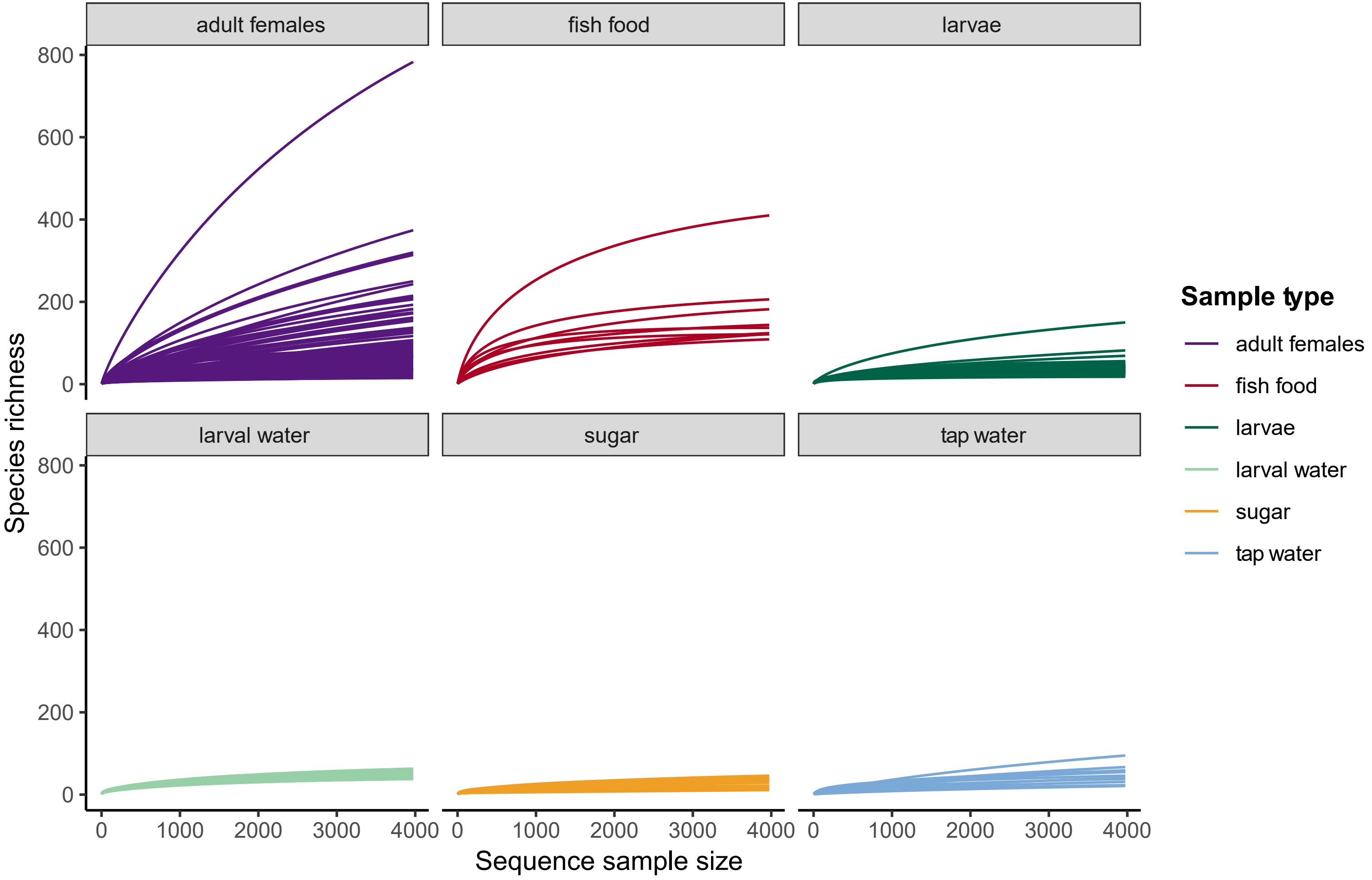

### Supp_fig_2.pdf

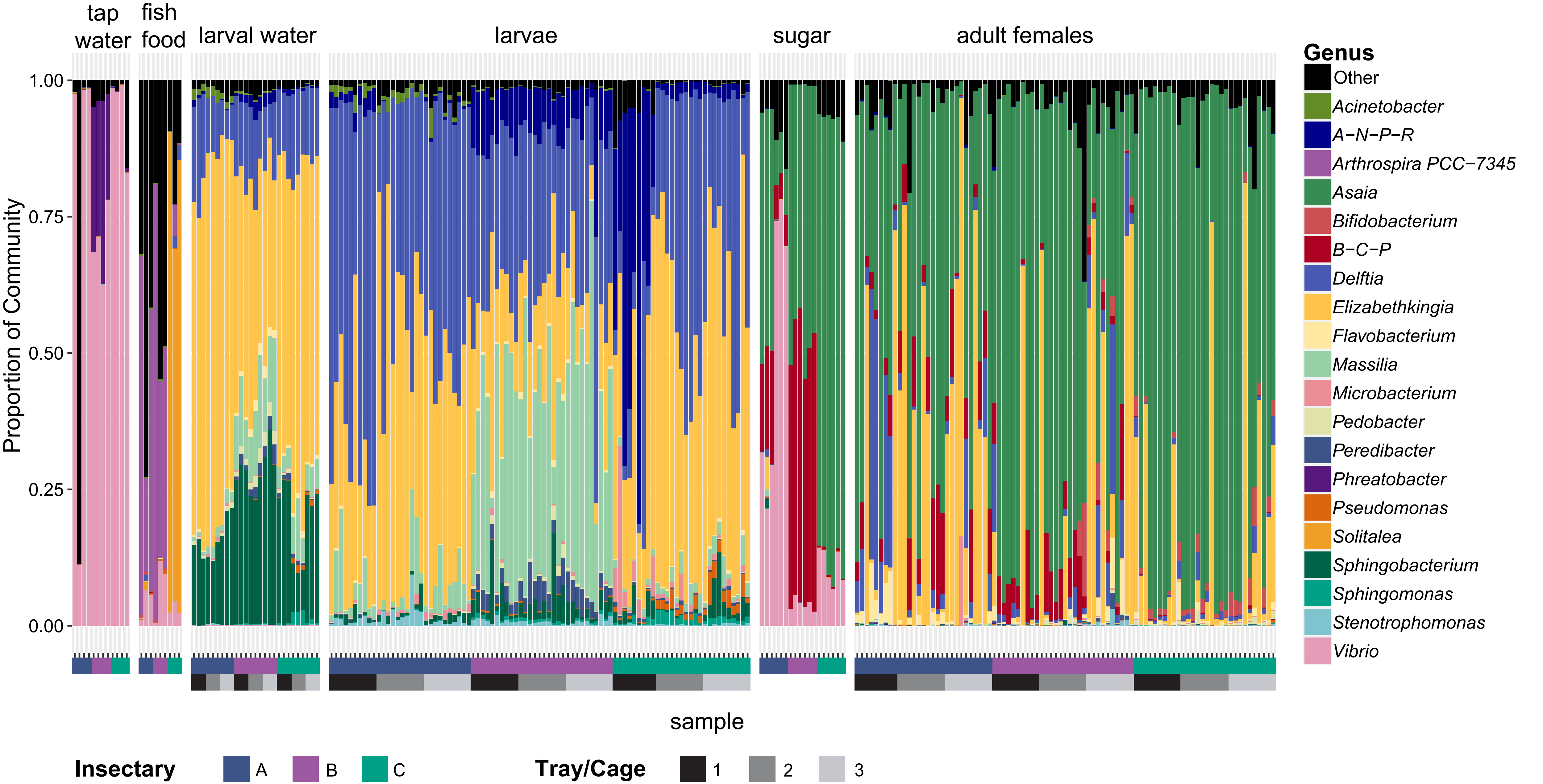

### Supp_fig_3.pdf

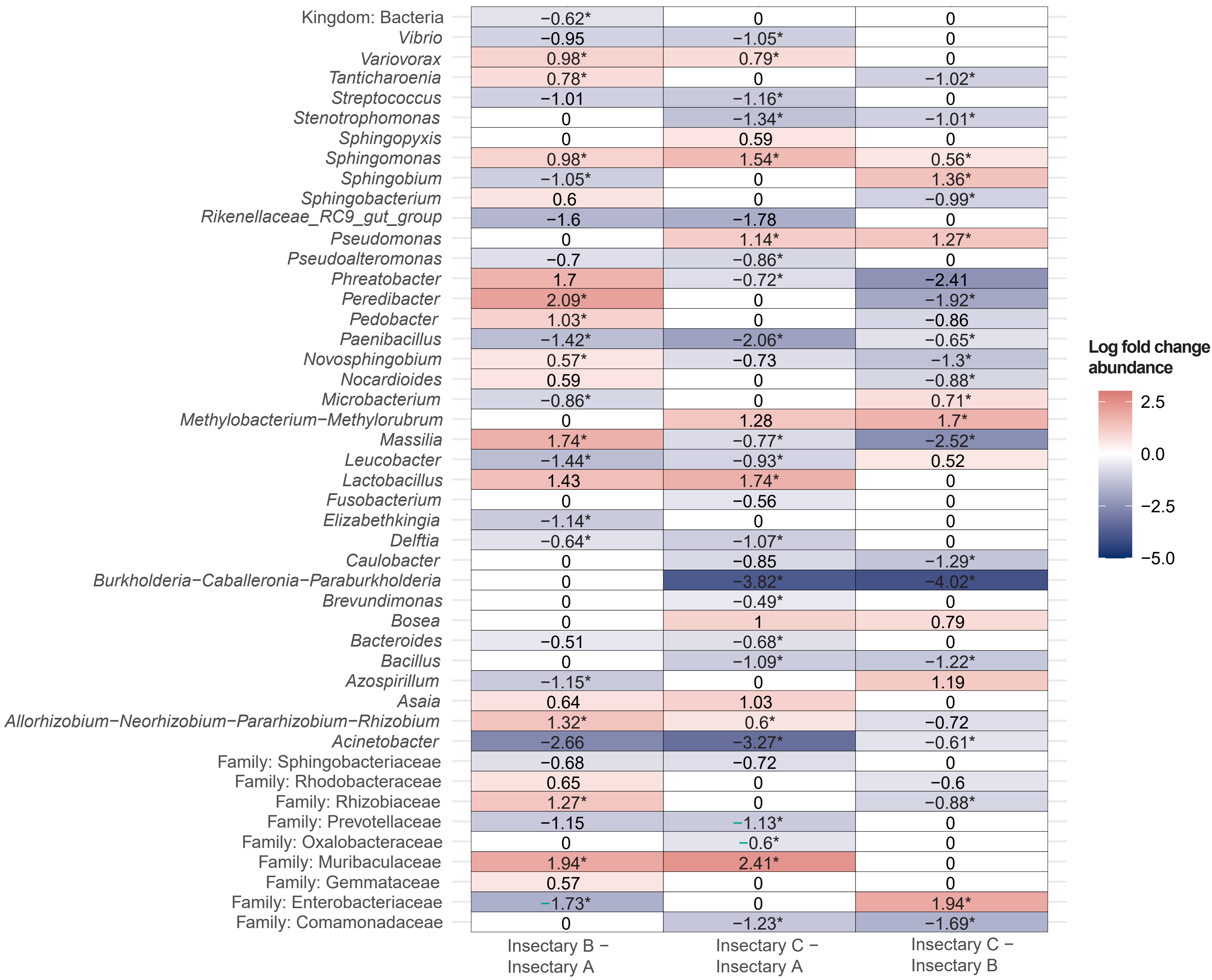

### Supp_fig_4.pdf

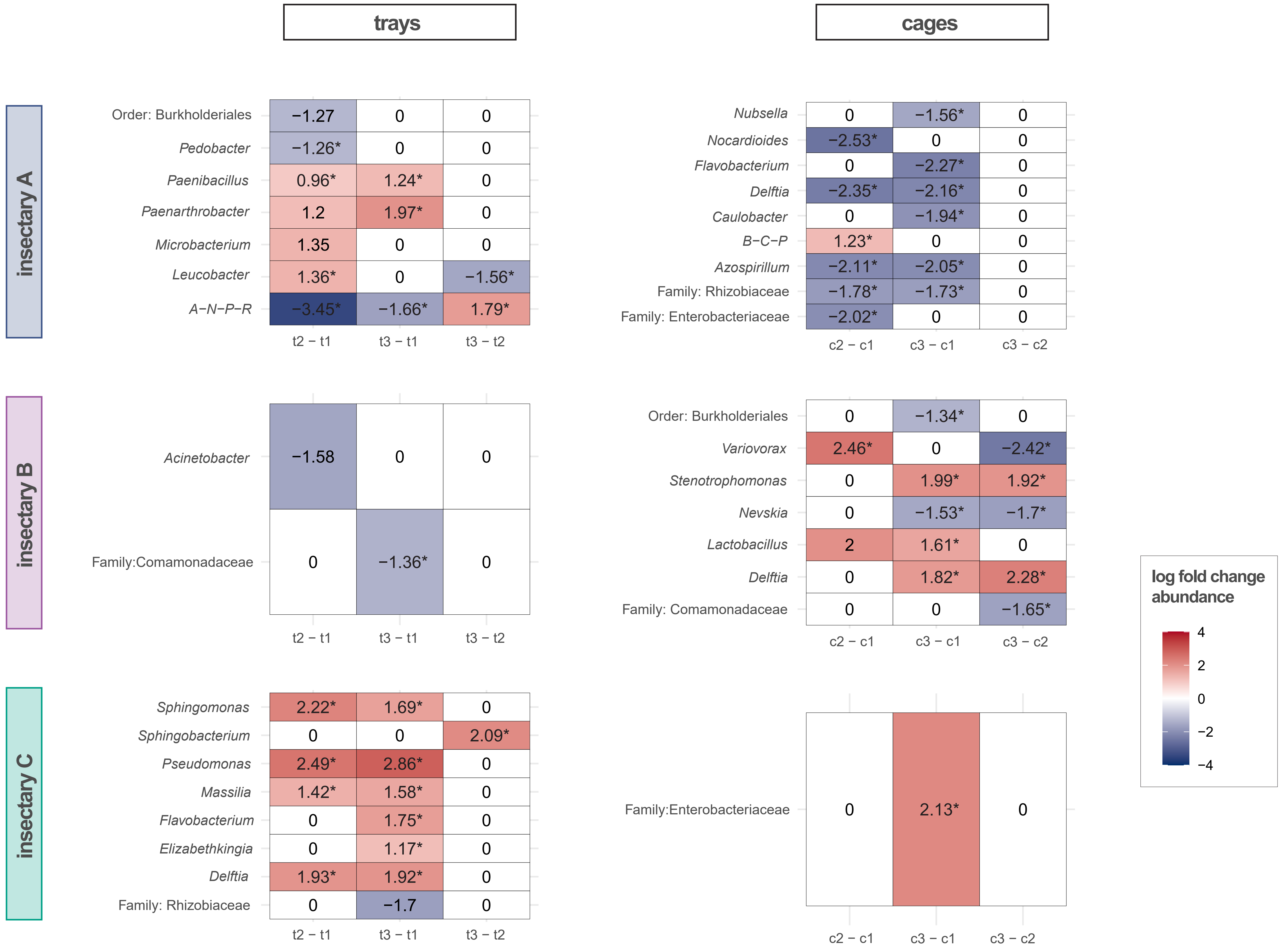
